# QstR-dependent regulation of natural competence and type VI secretion in *Vibrio cholerae*

**DOI:** 10.1101/337857

**Authors:** Milena Jaskólska, Sandrine Stutzmann, Candice Stoudmann, Melanie Blokesch

## Abstract

During growth on chitinous surfaces in its natural aquatic environment *Vibrio cholerae* develops natural competence for transformation and kills neighboring non-immune bacteria using a type VI secretion system (T6SS). Activation of these two phenotypes requires the chitin-induced regulator TfoX, but also integrates signals from quorum sensing via the intermediate regulator QstR, which belongs to the LuxR-type family of regulators. Here, we define the QstR regulon using RNA sequencing. Moreover, by mapping QstR binding sites using chromatin immunoprecipitation coupled with deep sequencing we demonstrate that QstR is likely a dual transcription factor that binds upstream of the up- and down-regulated genes. Like other LuxR-type family transcriptional regulators we show that QstR function is dependent on dimerization. However, in contrast to the well-studied LuxR-type biofilm regulator VpsT of *V. cholerae,* which requires the second messenger c-di-GMP, we show that QstR dimerization and function is c-di-GMP independent. Surprisingly, although ComEA, which is a periplasmic DNA-binding protein essential for transformation, is produced in a QstR-dependent manner, QstR-binding was not detected upstream of *comEA* suggesting the existence of a further regulatory pathway. Overall these results provide detailed insights into the function of a key regulator of natural competence and type VI secretion in *V. cholerae*.

## Introduction

*Vibrio cholerae,* the human pathogen responsible for the diarrheal disease cholera, is commonly found in aquatic environments, often in association with the chitinous exoskeletons of zooplankton and shellfish (1). Chitin not only provides a colonization surface and a source of nutrients, but also significantly influences the gene expression profile of *V. cholerae* (2) including induction of the physiological state of natural competence for transformation (3). Natural competence for transformation, the ability of a bacterium to actively take up DNA from the environment and maintain it in a heritable state resulting in transformation, is widespread throughout the bacteria with ˜ 80 species experimentally shown to be transformable. Moreover, competence development is often tightly regulated in response to specific environmental signals (4,5).

The regulatory network controlling competence development in *V. cholerae* is complex (recently reviewed in (6)) and is centered around the chitin-induced regulator TfoX, which in turn activates the expression of the genes encoding the DNA uptake machinery (3). In addition to chitin, competence development also requires other signals such as the unavailability of certain carbon sources (via <u>c</u>AMP <u>r</u>eceptor <u>p</u>rotein - CRP; (7)) or the regulatory protein CytR (8) and high cell density (via the regulator HapR; (9–11)).

High cell density, measured through the local concentration of secreted autoinducers (*Vibrio* species specific cholera autoinducer 1 [CAI-1] and autoinducer 2 [AI-2]), leads to the accumulation of the master regulator of quorum sensing (QS) HapR (12). HapR was shown to be crucial for full expression of *comEA* and *comEC* (3,9–11), which encode, respectively, a periplasmic DNA-binding protein required for DNA-uptake and the predicted inner membrane channel responsible for translocation of the incoming DNA into the cytoplasm (13–15). Additionally, HapR represses *dns,* which encodes an extracellular DNase that degrades transforming material (11,16).

We previously showed that HapR regulates *comEA* and *comEC* indirectly via an intermediate regulator, which we termed QstR (<u>QS</u> and <u>T</u>foX-dependent <u>R</u>egulator) (17). QstR is necessary for transformation and both TfoX and HapR regulate its production, thus, integrating both chitin and QS regulatory pathways. Furthermore, we showed that HapR directly binds to the *dns* promoter to repress it, but that further repression is achieved through QstR (17). However, the mechanism by which QstR regulates these genes is unknown.

Recently our group demonstrated that in addition to competence induction, growth on chitin leads to TfoX-dependent activation of the type VI secretion system (T6SS) (18). Co-regulation of competence with the T6SS, which forms a molecular killing device (19,20), allows the killing of neighboring non-immune cells followed by uptake of their DNA, and thus, promotes horizontal gene transfer on chitin surfaces (18). Notably, the TfoX mediated expression of T6SS genes is QS and QstR-dependent, as demonstrated by expression profiling, T6SS activity testing, and imaging on chitinous surfaces (18).

QstR is predicted to possess a LuxR-type C-terminal DNA-binding helix-turn-helix (HTH) domain (17). Proteins belonging to the LuxR-type superfamily are response regulators that often dimerize and bind DNA in response to phosphorylation (e.g. NarL of *Escherichia coli* and FixJ of *Sinorhizobium meliloti* (21,22)), autoinducers (e.g. LuxR of *Vibrio fischeri* and TraR of *Agrobacterium tumefaciens* (23,24)) or ligand binding (maltotriose and ATP for MalT of *E. coli* and c-di-GMP for VpsT of *V. cholerae* (25,26)). The C-terminal domain of QstR shares significant homology with the LuxR-type regulator VpsT, which can directly sense the secondary messenger c-di-GMP and controls biofilm matrix production in *V. cholerae.* The binding of c-di-GMP to VpsT elicits a conformational change that leads to dimer formation, which is required for DNA-binding and transcriptional activation of the target genes (26). The conserved c-di-GMP binding motif W[F/L/M][T/S]R of VpsT is present in QstR, however, with proline in the third position, raising the question of how this substitution might impact the activity of QstR.

Here, we define the QstR regulon and identify additional QstR regulated genes. We go on to assess the contribution of these genes to natural transformation and T6SS mediated killing, as well as the interplay between TfoX and QstR in their regulation. Additionally, we demonstrate that QstR binds to DNA directly upstream of the regulated genes and, similarly to VpsT, that dimerization is required for DNA-binding. In contrast to VpsT however, elevated intracellular c-di-GMP concentrations are not required for QstR activity. Furthermore, we provide evidence that QstR-dependent activation of *comEA* expression might be achieved via an as yet unidentified intermediate regulator.

## Materials and Methods

### Bacterial strains, plasmids and growth conditions

Bacterial strains and plasmids used in this study are listed in Table S1. *E coli* strains DH5α, TOP10, XL10-Gold were used for cloning and derivatives of strains S17-1λpir and MFDp/r served as donors in mating experiments with *V. cholerae.* Bacteria were cultured in LB medium or on LB Agar (Luria/Miller; Carl Roth) at 30°C or 37°C as required. Where appropriate the following antibiotics were used for selection: ampicillin (Amp, 50 or 100 μg/ml), kanamycin (Kan, 75 μg/ml for *V. cholerae;* 50 μg/ml for *E. coli*), chloramphenicol (Cm, 2.5 μg/ml), gentamicin (Gent, 50 μg/ml), streptomycin (Str, 100 μg/ml) and rifampicin (Rif, 100 μg/ml). Thiosulfate citrate bile salts sucrose agar (TCBS; Sigma-Aldrich) supplemented with appropriate antibiotics was used for selection of *V. cholerae* following mating with *E. coli*. For strain construction or to assess natural transformability on chitin bacteria were grown on chitin flakes in 0.5 x defined artificial sea water (DASW) supplemented with 50 mM HEPES and vitamins (MEM, Gibco) as previously described (3,27). Where indicated the growth medium was supplemented with 0.02% or 0.2% arabinose to induce expression from the P_BAD_ promoter. To visualize β-galactosidase (LacZ) activity plates were supplemented with 5-bromo-4-chloro-3-indolyl-β-D-galactopyranoside (X-gal, AppliChem) at a final concentration 40 μg/ml. To induce expression from the bacterial two-hybrid plasmids 0.5 mM Isopropyl β-D-1-thiogalactopyranoside (IPTG, AppliChem) was used. The counter-selection following biparental mating with *E. coli* carrying pGP704-Sac28 derivatives was done on NaCl-free LB plates supplemented with 10% sucrose.

### Strain construction

DNA manipulations and molecular cloning were performed according to standard procedures (28). Genetic engineering of *V. cholerae* was carried out by the previously described TransFLP method (27,29) or using derivatives of counter-selectable plasmid pGP704-Sac28 delivered by biparental mating from *E. coli* (2). To insert the *itfoX* fragment *(araC* plus *tfoX* under the control of P_BAD_ promoter) in the *lacZ* gene a *pheS*\*-based counter selection was used ((30), van der Henst *et al.,* BioRxiv: https://doi.org/10.1101/235598). All constructs were verified by colony PCR and sequencing (Microsynth, Switzerland). The mini-Tn7 transposon carrying *araC* and either *tfoX, qstR,* qstR[L137A], *vdcA* or *cdpA* under control of P_BAD_ promoter was delivered by triparental mating as described previously (31) and the insertion in the *V. cholerae* chromosome was confirmed by PCR.

### Sequence alignment

QstR (VC0396, NP_230050) and VpsT (VCA0952, NP_233336) protein sequence alignment was generated with Clustal Omega (32) and edited with ESPript 3.0 (33).

### Transformation frequency assay

Transformability of *V. cholerae* strains was assessed either on chitin as previously described (27,29) or in chitin-independent transformation assays with expression of an arabinose-inducible chromosomal copy of *tfoX* either from *TntfoX* (11) or *itfoX*. All strains were grown to a similar OD_600_ of ˜1.4. Genomic DNA of a strain carrying a kanamycin resistance cassette in the *lacZ* gene was used as transforming material. Transformation frequencies were calculated as the ratio of kanamycin resistant transformants to the total number of bacteria.

### Interbacterial killing assay

The interbacterial killing assay was performed according to the previously published protocol (18). Briefly, *V. cholerae* strains were grown to OD_600_ ˜ 1.5 in the culture medium without or with 0.2% arabinose to induce *tfoX* and/or *qstR* from the transposon and were mixed with *E. coli* TOP10 strain in 10:1 ratio (*V. cholerae* to *E. coli).* 5 μl of the mixture was spotted on sterile PES membrane filters placed on prewarmed LB plates (± 0.2% arabinose) and incubated for 4 h at 37°C. The bacteria were subsequently washed from the filters and serial dilutions were spotted on LB plates supplemented with streptomycin to select for surviving prey cells (e.g., streptomycin-resistant *E. coli*).

### Western blotting

The cultures for Western blotting were grown in the indicated medium (± inducer), harvested, and resuspended in an appropriate volume of 2 x Laemmli buffer (Sigma-Aldrich) according to the to optical density at 600nm of the initial culture (OD_600_) followed by 15 min incubation at 95°C. The proteins were resolved by SDS-PAGE (15%) and transferred to a PVDF membrane (Merck Millipore) using a wet-or semidry transfer apparatus. After incubation with the blocking buffer (2.5% skimmed milk in 1 x Tris-buffered saline with 0.1% Tween-20), one of the following primary antibodies was added: anti-QstR (raised in rabbits against synthetic peptides; Eurogentec #1412414) at 1:2,500 dilution; anti-ComEA (raised in rabbits against synthetic peptides; Eurogentec #GP1248 (34)), anti-Hcp (against synthetic peptide, Eurogentec #1510528 (35)), anti-HapR (against synthetic peptide, Biomatik #A000542 (11)), all at 1:5,000 dilution; anti-RNAP-beta (Neoclone #WP001; raised in mouse) at dilution 1:2,000. The secondary antibodies used were anti-rabbit IgG HRP (Sigma-Aldrich #A9169) and anti-mouse IgG HRP (Sigma-Aldrich #A5278), both at 1:10,000 dilutions. The proteins were visualized by chemiluminescence using Lumi-Light^PLUS^ Western Blotting Substrate (Roche).

### β-galactosidase assay

Overnight culture of each strain was spotted on LB plates containing 100 μg/ml ampicillin, 50 μg/ml kanamycin, 0.5 mM IPTG and 40 μg/ml X-gal. After 24 h incubation at 30°C the cells were collected, resuspended in sterile phosphate buffered saline (PBS) and the suspensions were diluted 10-fold. 500 μl of cell suspension was used in the assay and diluted if necessary. β-galactosidase activity was then measured using the previously established protocol (36).

### Protein purification

N-terminally tagged (Strep-tagII) wild-type QstR and QstR[L137A] were produced from pBAD-qstR-N-strep and pBAD-qstR-N-strep[L137A] plasmids, respectively, in *E. coli* strain BL21 (DE3) (Agilent). The strains were grown in LB medium with shaking at 30°C to OD_600_ ˜ 0.8, then arabinose at a final concentration of 0.2% was added and the cultures were further incubated at 16°C overnight. The harvested cell pellets were resuspended in lysis buffer (50 mM sodium phosphate pH 8.0, 125 mM NaCl, 1% Triton) supplemented with cOmplete Mini EDTA-free protease inhibitor cocktail and DNaseI at a final concentration of 20 μg/ml (both Roche). The cells were lysed by repeated passing through a French press cell and the lysates were cleared by centrifugation at 17000 rpm for 30 min at 4°C. The cleared lysates were applied to columns packed with Strep-Tactin^®^ Sepharose (IBA, Germany) equilibrated with Buffer W (100 mM Tris-HCl pH 8.0, 150 mM NaCl, 1 mM EDTA) and allowed to pass through by gravity flow. The columns were washed 5 times with two column volumes of the washing buffer (Buffer W with 500 mM NaCl). The strep-tagged proteins were eluted in six fractions with buffer containing 100 mM Tris-HCl pH 8.0, 300 mM NaCl, 1 mM EDTA and 2.5 mM D-desthiobiotin. The fractions with highest amounts of proteins, as estimated by Coomassie Brilliant Blue staining after SDS PAGE (15%) separation were pooled and concentrated using Amicon Ultra 10K centrifugal filter units (Merck Millipore). The protein concentration was determined using Bradford Reagent (Sigma-Aldrich).

### Analytical size exclusion chromatography (SEC)

Purified wild-type QstR or QstR[L137A] proteins were applied to Superdex 200 10/300 GL (GE Healthcare) gel filtration column. Samples were run in buffer containing 100 mM Tris-HCl pH 8.0, 300 mM NaCl and 1 mM EDTA at the flow rate 0.5 ml/min. The standard curve was obtained by applying a set of known protein standards (conalbumin 75000 Da, ovalbumin 43000 Da, carbonic anhydrase 29000 Da, ribonuclease A 13700 Da) to the column and plotting partition coefficients (K_av_, calculated according to manufacturer’s instructions) against Log10 of their molecular weight. The molecular weight of QstR and QstR[L137A] was estimated by comparing their partition coefficients to the standard curve.

### Quantitative Reverse Transcription PCR (qRT-PCR)

Bacterial growth, RNA purification, cDNA synthesis and quantitative PCR were performed as previously described (11). LightCycler Nano or LightCycler 96 systems (Roche) were used for qPCR runs. Expression values in the graphs are presented as relative to the mRNA levels of the reference gene *gyrA.*

### RNA sequencing (RNA-seq)

Bacterial growth and RNA purification (three replicates per sample) was done as previously described (18). Further processing of the samples and data analysis was conducted by Microsynth (Balgach, Switzerland) using Illumina RNA Sequencing for Differential Gene Expression Analysis pipeline as described in (18).

### Chromatin Immunoprecipitation (ChIP)

ChIP was performed as described in (37) with modifications. 10 ml (for ChIP-qPCR) or 100 ml (for ChIP-seq) cultures were grown in LB medium supplemented with 0.02% arabinose unless stated otherwise. The cells were transferred to Falcon tubes and cross-linked with formaldehyde added to the final concentration of 1% for 10 min at room temperature followed by 30 min incubation on ice. After two washes with sterile PBS the pellets were resuspended in TES buffer (10 mM Tris-HCl pH 7.5, 100 mM NaCl, 1 mM EDTA) and bacteria lysed with Ready-Lyse Lysozyme Solution (Epicentre) in presence of protease inhibitors (Roche). The lysates were then sonicated to shear DNA (3 × 10 cycles; 30s on, 30s off per cycle; Diagenode) and centrifuged at max speed to remove cell debris. The supernatants were adjusted to 1 ml with ChIP buffer (16.7 mM Tris-HCl pH 8.1, 167 mM NaCl, 1.2 mM EDTA, 1.1% Triton X-100, 0.01% SDS). For ChIP-qPCR ˜ 4 mg of total protein was used per sample, for ChIP-seq the obtained lysate was split in two and filled up to 1 ml with ChIP buffer. The diluted lysates were pre-cleared with Protein A Dynabeads (Life Technologies) before being incubated overnight with the anti-QstR antibody. The immuno-complexes were captured by incubation with Protein A Dynabeads, washed and eluted as described (37). The cross-link was reversed by incubation at 65°C overnight and the DNA was isolated using phenol:chloroform:isoamyl alcohol (25:24:1). For ChIP-qPCR 10% of each sample was removed after the pre-clearing step and the extracted DNA served as input. 1 μl of ChIP samples and diluted input samples served as templates for qPCR performed with primers annealing upstream of the QstR-regulted genes and upstream of *gyrA* as control. Input % was calculated as follows: Input % = 100/2^ΔCq[normalised ChIP] where ΔCq[normalised ChIP]=Cq[ChIP]-(Cq[input]-log2[input dilution factor]). To calculate fold enrichment, the ‘Input %’ for the target gene was divided by ‘Input %’ for *gyrA* in the same sample.

For the ChIP-seq samples (two replicates per sample), the library preparation, sequencing (Illumina NextSeq 500, mid-output, v2, 2×75bp), and data analysis was carried out by Microsynth (Balgach, Switzerland). The reads were mapped to the reference sequence of *V. cholerae* O1 El Tor strain N16961 (accession numbers NC_002505 and NC_002506 for both chromosomes).

### Transposon mutagenesis screen

For generation of transposon mutants libraries, strain A1552Δ*lacZ-comEA-lacZ::FRT-TntfoX* was mated with *E. coli* strain MFDpir carrying pSC189 plasmid that contains the *mariner-based* transposon (38). The strains were co-cultured on LB agar plates containing diaminopimelic acid (DAP 0.3 mM, required for growth of MFDpir strain (39)). After 6 h at 37°C the cells were collected, resuspended in sterile PBS, and plated on LB plates containing kanamycin and X-gal. Two screens were performed, one with ˜20,000 colonies and one with ˜ 60,000 colonies inspected. To identify the site of transposon insertions in the purified clones two-step arbitrary PCR adapted from (40) with GoTaq polymerase (Promega) was performed. In round 1, primers ARB6/ARB7 and MarTransp-Kan-1 (5’ CTT CCT CGT GCT TTA CGG TAT CGC) specific to the kanamycin resistance cassette in pSC189 were used (95°C 10min; 6 × 95°C 15 s, 30°C 30 s, 72°C 1min 15 s; 30 × 95°C 15 s, 52°C 30 s, 72°C 1min 15 s; 72°C 5 min). For round 2, primers ARB2 and MarTransp-Kan-2 (5’ TTC TGA GCG GGA CTC TGG GGT ACG) were used (95°C 2 min; 32 × 95°C 15 s, 52°C 30 s, 72°C 1 min 15 s; 72°C 5 min) with purified PCR product from round 1 as template. Purified PCR products from round 2 were sequenced with MarTransp-Kan-2 primer (Microsynth, Switzerland) and aligned to the genomic DNA sequence.

### DNA affinity pull-down assay

30 ml cultures of strain A1552-*TntfoX* and A1552Δ*qstR-TntfoX* were grown in the presence of 0.2% arabinose and harvested at OD_600_ ˜2. The lysis, probe binding, incubation with the lysate and elution were performed as described in (41). The biotinylated probes contained the fragment upstream of *comEA* (−341 and +74 relative to the start codon), *qstR* (−337 and +78 relative to the start codon) and *gyrA* (−360 and +75 relative to start codon). The fractions eluted at 500 mM NaCl were loaded on 15% SDS polyacrylamide gel and subjected to short migration (until the dye reached ˜ 2 cm from the well). The lanes were excised and proteins identified by liquid chromatography-tandem mass spectrometry (LC-MS/MS, EPFL Proteomics Core Facility).

## Results

### Mutation in the putative dimerization interface abolishes the functionality of QstR

The C-terminal domain of QstR shares homology (35% amino acids identity and 61% similarity) with the well-studied biofilm regulator VpsT (Fig. 1A). VpsT undergoes c-di-GMP dependent dimerization, which is required for DNA binding and transcriptional activation of the target genes (26). QstR contains a variant of the conserved c-di-GMP binding motif found in VpsT, with a proline instead of a threonine in the third position (WLPR instead of WLTR). Given the properties of proline this substitution is likely to affect the ability of QstR to bind c-di-GMP. Therefore, to investigate how QstR regulates the known subset of competence and T6SS genes, we introduced mutations corresponding to areas that encode VpsT’s c-di-GMP binding motif and c-di-GMP dependent dimerization interface into the native *qstR* gene (Fig. 1A). We then assessed the transformability of the strains producing these QstR variants in a chitin-independent transformation assay (using an integrated transposon harboring an arabinose-inducible copy of *tfoX*(*TntfoX*) to induce competence, as previously established (11)).

**Figure 1.**
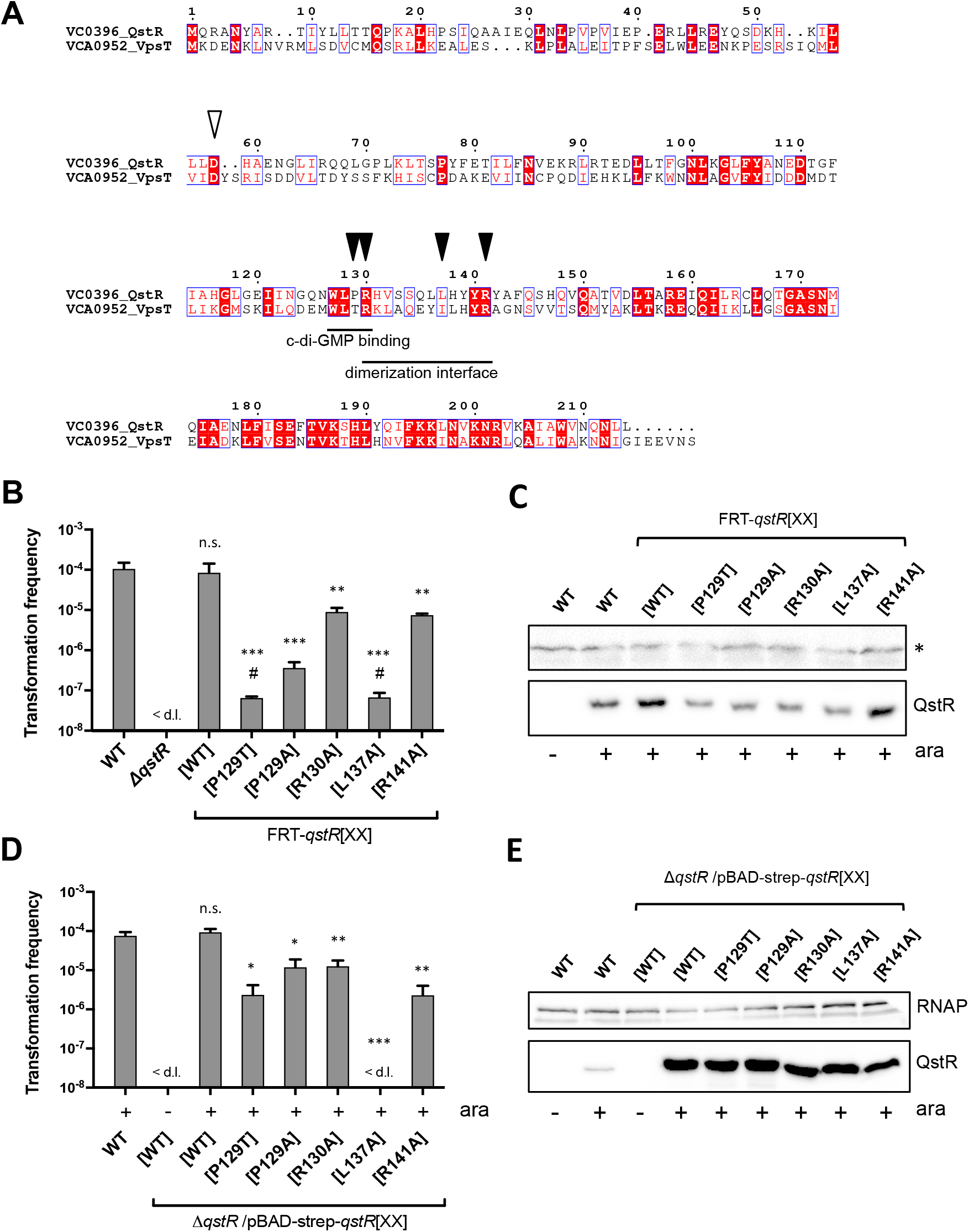
Functionality of QstR variants. **(A)** Alignment of the protein sequence of QstR and VpsT. The c-di-GMP motif and the dimerization interface of VpsT are highlighted and the residues replaced in QstR to create mutants are marked with closed arrows. The putative phosphorylation site is marked with an open arrow. **(B)** and **(D)** Transformation frequencies of strains encoding wild-type QstR and five QstR variants carried either at *qstR’s* native chromosomal locus (B) or on a plasmid to test complementation of a *qstR* deletion strain (D). All strains carry an inducible copy of *tfoX (TntfoX)* and were grown with 0.02% arabinose to induce *tfoX* and therefore competence. In (D) arabinose also induced the expression of *qstR* variants from the plasmid where indicated. Strains marked as FRT-qstR[XX] contain a FRT scar upstream of the *qstR* gene (see main text). The values shown are averages of three independent experiments with error bars representing the standard deviation (SD). < d.l., below detection limit; #, transformation frequency below the detection limit in one ([P129T]) or two ([L137A]) experiments. When the transformation frequency was below the detection limit, the limit of detection was used to calculate the average. Significant differences between WT and the strains producing QstR variants were determined by Student’s t-test (* p<0.05, ** p<0.01, *** p<0.001; n.s., not significant). For transformation below the detection limit the latter was used for statistical analyses. **(C)** and **(E)** QstR levels produced by strains used in (B) and (D) were assessed by Western blotting with anti-QstR antibody. (−) and (+) indicates absence or presence of 0.02% arabinose (ara) in the culture medium. A non-specific cross-reaction band (* in C) and the anti-RNAP band (E) served as loading controls.

Since the genetic engineering method (TransFLP) used to introduce mutations leaves an FRT scar upstream of the *qstR* promoter, a strain with an FRT scar upstream of wild-type *qstR* (hereafter referred to as FRT-*qstR*) was included in our analysis as control. This strain was previously shown to exhibit wild-type levels of competence induction and transformation (34). As shown in Fig. 1B restoring the c-di-GMP binding motif of VpsT (FRT-*qstR*[P129T]) significantly affected the ability of QstR to support transformation, with frequencies below or at the limit of detection. Changing proline to alanine also resulted in significantly reduced transformation frequencies (˜ 100-fold). The arginine in the c-di-GMP binding motif WLTR, which is conserved among the VpsT homologs, was also shown to be essential for VpsT function with a R134A mutant unable to complement a *vpsT* deletion (26). However, replacing the corresponding residue of QstR (i.e. R130A) had only a modest effect, with a 10-fold reduction in transformation. This result demonstrates that this residue is not critical for QstR function. Moreover, since R134 of VpsT is responsible for the interaction with c-di-GMP, it suggests that QstR might not interact with c-di-GMP or, at least, not in a manner similar to VpsT.

A VpsT variant with the hydrophobic residue in the c-di-GMP dependent dimerization interface replaced by a polar amino acid (I141E) is not functional, due to an inability to form c-di-GMP stabilized dimers (26). Replacing the analogous residue in QstR (FRT-*qstR*[L137A]) resulted in the detection of only rare transformation events (Fig. 1B), suggesting that QstR, like VpsT, might require oligomerization in order to activate the target genes. In contrast, changing the conserved arginine at the edge of the dimerization interface (26) resulting in QstR[R141A] had only a minor effect on transformation (Fig. 1B).

Since Western blotting indicated that most of the QstR variants were produced at reduced levels (Fig. 1C) we repeated the transformation experiments in a *qstR* deletion background, with each QstR variant overproduced from a plasmid. Even though the protein levels of the QstR variants were now considerably higher than that of the chromosomally encoded QstR (Fig. 1E), the transformation results were similar for the [R130A] and [R141A] variants, and although the transformation frequencies for the variants with proline substitutions were markedly increased, they remained significantly lower than for WT QstR (Fig. 1D). Importantly, even when overproduced, the QstR[L137A] variant was unable to complement the *qstR* deletion (Fig. 1D) highlighting the importance of this residue for QstR function.

The activity of many members of the LuxR-type family of regulators is affected by their phosphorylation state (42). Thus, we introduced substitutions into the putative phosphorylation site of QstR (Fig. 1A), which is shared with VpsT and other VpsT homologs in *Vibrio* species (26), designed to either abolish phosphorylation (D58A) or mimic a constitutively active state (D58E). Similar to VpsT (26), these variants did not affect QstR activity as assessed by transformation (Fig. S1), suggesting that phosphorylation does not regulate QstR activity.

### Dimer formation of QstR is required for its functionality

To further examine whether QstR is capable of oligomerization we employed a bacterial two-hybrid system, based on reconstitution of the catalytic domain of adenylate cyclase (43). Thus, to investigate QstR self-interaction we fused QstR to either the N- or C-terminus of the T18 and T25 adenylate cyclase domains and transformed *E. coli* strain BTH101 with all possible plasmid combinations. Indeed, we observed an interaction between QstR in all combinations, as evidenced by blue color development (Fig. S2A). Next, we inspected the interactions between the QstR variants described above (Fig. 1B). We observed a self-interaction for QstR[R130A] and [R141A], although it was reduced as compared to wild-type QstR. In contrast, no interaction was detected for the [P129T], [P129A] and [L137A] variants (Fig. 2A), consistent with the functional analysis of these mutants (Fig. 1B). We also analyzed the ability of wild-type QstR to interact with the mutant variants and detected an interaction with QstR[P129T] and [P129A] but not QstR[L137A] (Fig. S2B), indicating that a substitution in this position abolishes any QstR-QstR interaction.

**Figure 2.**
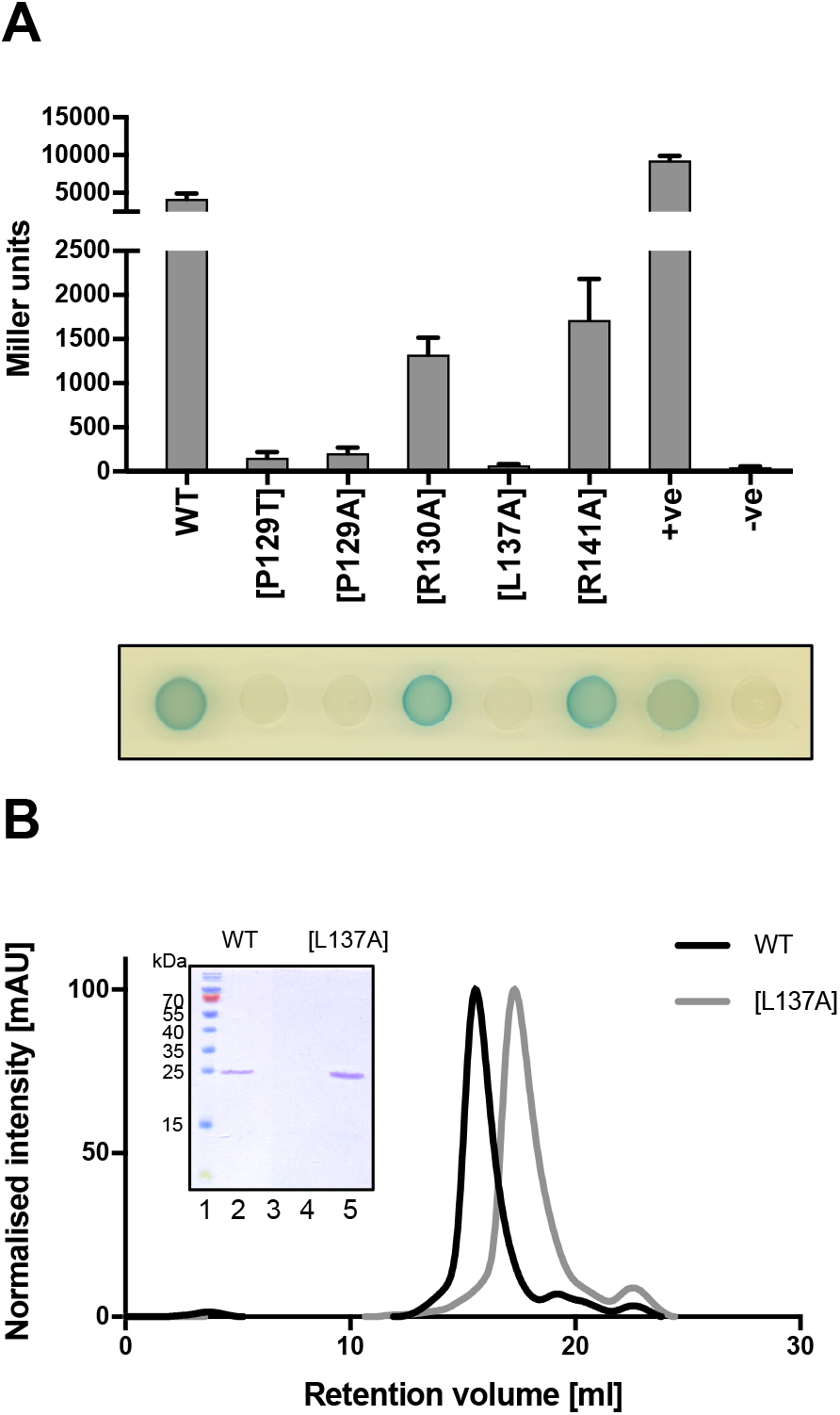
QstR forms dimers. **(A)** Bacterial two-hybrid system (BACTH) to investigate the interaction of wild-type (WT) QstR and between QstR mutants. WT and variants of QstR were fused to the T18 and T25 fragments of adenylate cyclase and the interactions were examined in the *E. coli* BTH101 strain. The overnight cultures of the strains carrying the *qstR* fusions on the BACTH plasmids were spotted on two-hybrid LB plates and scanned after 24 h incubation at 30°C. To measure β-galactosidase activity the cells were scraped from the plate and resuspended in PBS. The strains carrying pKT25-zip/pUT18C-zip and pUT18/pKNT25*qstR* served as positive (+ve) and negative (-ve) controls, respectively. The values are averages of three independent experiments with error bars representing the SD. **(B)** Size exclusion chromatography of QstR and the QstR[L137A] variant. Affinity purified strep-tagged QstR and QstR[L137A] were applied to a Superdex 200 10/300 GL gel filtration column. The inset shows the SDS-PAGE analysis of the concentrated peak fractions: lane 1, pre-stained protein ladder; lane 2, concentrated peak fractions of WT QstR (eluted at ˜ 15 ml); lane 3, concentrated fraction of the WT QstR eluted at ˜ 22 ml; lane 4, empty; lane 5, concentrated fraction of the QstR[L137A] variant (eluted at ˜ 17 ml).

Next, we purified a fully functional strep-tagged version of QstR (Fig. 1D) and QstR[L137A] and performed analytical size exclusion chromatography (SEC). As shown in Fig. 2B, we observed a shift in the protein peak elution volume between wild-type QstR and QstR[L137A]. This was not due to QstR[L137A] degradation, as evidenced by SDS-PAGE separation and Coomassie Brilliant Blue staining of the concentrated elution fractions (Fig. 2B inset). The calculated size of QstR was approximately double that of QstR[L137A] (Fig. S3), consistent with wild-type QstR behaving as a dimer and QstR[L137A] as a monomer. Collectively, these data suggest that QstR forms dimers via the dimerization interface, with a crucial role for residue L137 in this interaction. Furthermore, given the inability of QstR[L137A] to support transformation, dimerization seems to be essential for QstR function.

### QstR is functional under low c-di-GMP concentration

The ability of the QstR[R130A] variant to support transformation and the observation that restoring the canonical c-di-GMP binding motif significantly impaired QstR function, hinted that QstR might not interact with c-di-GMP in the same manner as VpsT. Therefore, we decided to directly examine the effect of intracellular c-di-GMP on QstR’s function in a transformation assay. In order to modulate the intracellular levels of c-di-GMP we induced either *vdcA* (encoding a diguanylate cyclase that synthesizes c-di-GMP) or *cdpA* (encoding a phosphodiesterase that degrades c-di-GMP). Both enzymes have previously been used to successfully increase/decrease intracellular c-di-GMP levels (35,44). In the chitin-independent transformation assay we observed a small reduction in the transformation frequency (˜2-3-fold) when c-di-GMP concentration was reduced, for both WT and the FRT-*qstR* control strain (Fig. 3A). A similar effect was also observed on chitin (Fig. 3B). Nevertheless, this minor reduction of natural transformability was not statistically significant for all strains in which the c-di-GMP level was lowered. Indeed, these strains were still highly transformable under the tested conditions.

**Figure 3.**
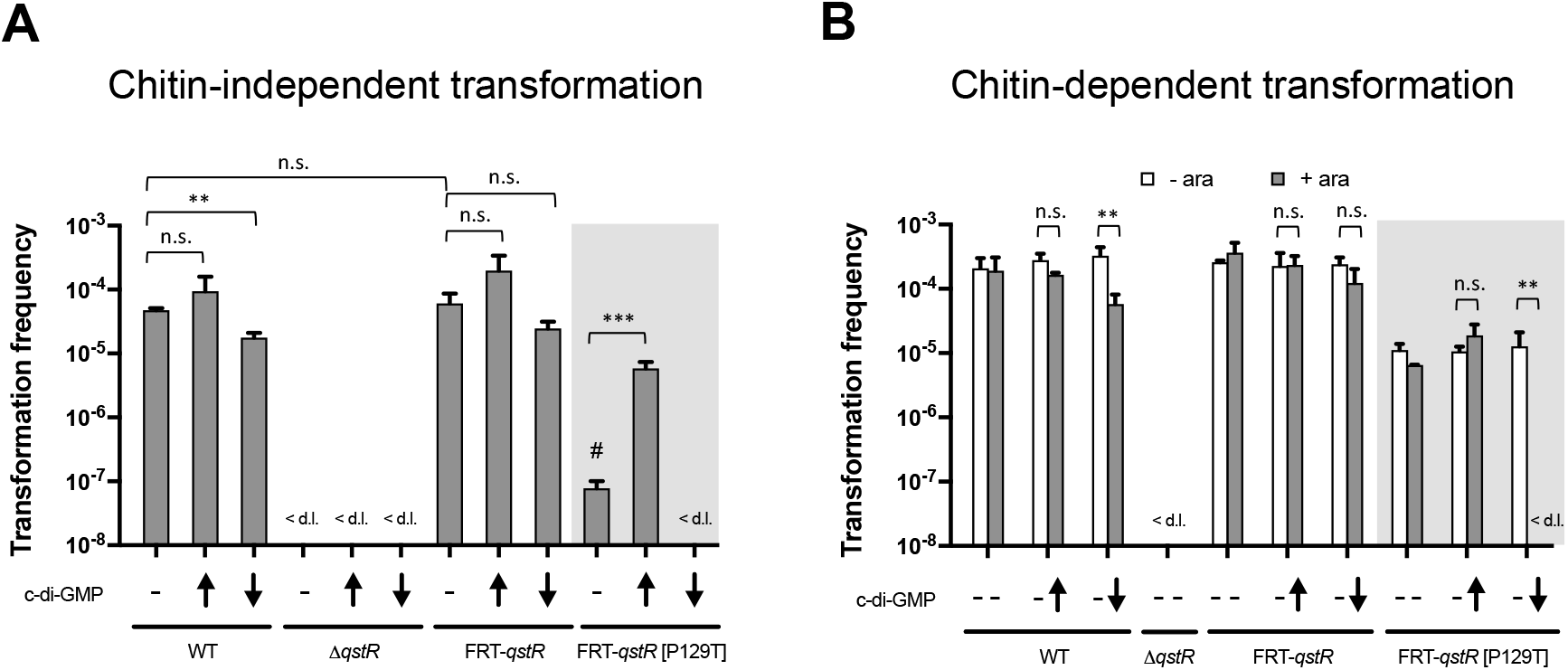
QstR is functional under low intracellular c-di-GMP concentrations while QstR[P129T] is c-di-GMP dependent. Transformation frequencies of strains carrying a chromosomal, arabinose-inducible copy of either *vdcA* or *cdpA* on the transposon to increase (↑) or decrease (↓) intracellular c-di-GMP levels, respectively, compared to isogenic strains without the transposon. **(A)** Transformation frequencies were determined in a chitin-independent transformation assay with all strains encoding an arabinose-inducible copy of *tfoX* at a neutral chromosomal locus (itfoX). The strains were grown with 0.2% arabinose to induce expression of *tfoX* and *vdcA/cdpA*. **(B)** Transformation was assessed upon growth on chitin. Where indicated, 0.2% arabinose was added to induce *vdcA/cdpA* expression. Due to the FRT scar upstream of *qstR* in FRT-*qstR*[P129T] (shaded box) the strains with a FRT scar upstream of *qstR* were used as additional control. The values are averages of three independent experiments with error bars representing the SD. <d.l., below detection limit; #, transformation frequency below the detection limit in two out of three experiments, thus the detection limit was used for calculation of the average for this strain. Significant differences were determined by Student’s t-test (* p<0.05, ** p<0.01, *** p<0.001; n.s., not significant). For undetectable transformation the detection limit was used for statistical analyses.

In contrast, in the chitin-independent assay the QstR[P129T] variant only became functional when c-di-GMP concentrations were increased (Fig. 3A). Remarkably, when grown on the natural competence inducer chitin, a strain producing this QstR variant was transformable without altering c-di-GMP levels, and artificially increasing the c-di-GMP levels did not significantly increase its transformability (Fig. 3B). Lowering c-di-GMP concentration, however, resulted in no detectable transformation events, suggesting that intracellular c-di-GMP concentrations are considerably higher during growth on chitinous surfaces compared to growth in LB medium. Taken together, the gain of function of QstR[P129T] under increased c-di-GMP levels in the chitin-independent assay and the loss of function under low c-di-GMP levels during growth on chitin, shows that this variant is strictly c-di-GMP dependent and demonstrates the effectiveness of *vdcA/cdpA* expression in altering intracellular c-di-GMP levels. Furthermore, since in both assays wild-type QstR was functional under low c-di-GMP conditions, these results suggest that this second messenger is not essential for QstR function.

### Defining the QstR regulon

We previously showed that QstR is required for the TfoX-dependent induction of *comEA, comEC* and T6SS genes, and that it also acts in concert with HapR to fully repress the extracellular nuclease *dns* (17,18). Here, to better understand QstR-dependent regulation of competence and T6SS we attempted to characterize the QstR regulon. Thus, we first analyzed a previously obtained RNA-seq data set (accession number GSE80217, for strain A1552 GSE79467) comparing the expression profiles between strain A1552 (WT), A1552 carrying an inducible copy of *tfoX* within a transposon (A1552-Tn*tfoX*) and the isogenic strains with either *hapR* (required for *qstR* expression) or *qstR* deleted (18,35).

In addition to the above-mentioned genes, several other genes that were induced at least 2-fold by TfoX also required QstR for full induction. These genes encode proteins involved in natural competence such as *comM* and *comF,* as well as genes not known to affect either of the QstR-dependent phenotypes (competence or type VI secretion) such as *ligA2, VC0033* (next to *VC0032=comM*), *VC0542* and *VC1479* (Fig. 4A; a full list of TfoX-induced genes is provided in Supplementary file 1). As shown previously, *dns* repression by TfoX was abolished in Δ*qstR* strain, while the transcript levels were even higher in the *hapR* negative strain (Fig. 4A, shaded box) consistent with its role as the primary *dns* repressor (17). Notably, we also observed TfoX-dependent transcriptional repression of *tfoY,* which encodes the second master regulator of T6SS (35). Notably, this repression was absent or, at least, occurred to a lower extent, in the *qstR* and *hapR* mutants (Fig. 4A, shaded box).

To investigate whether QstR alone is sufficient to regulate the expression of the QstR-dependent genes, we inserted an arabinose-inducible copy of *qstR* into a mini-Tn7 transposon (analogous to *TntfoX* (11)), further referred to as TnqstR) and inserted this transposon into the WT strain. Next, we performed RNA-seq to determine the effects of QstR on the transcriptome compared to TfoX-dependent induction (in this experiment a fully functional strep-tagged version of *tfoX* was used; (35)). In addition, to evaluate whether HapR has any further role in the regulation of QstR-dependent genes, other than *qstR* activation, we included a Δ*hapR-TnqstR* strain in the analysis. This strain also carried a *vpsA* deletion to counteract the increased biofilm production by the Δ*hapR* mutant. Overall, 51 genes were induced at least 2-fold when QstR alone was produced from the transposon, including all the genes identified as QstR-dependent in the previous analysis (Supplementary file 2). Several of the QstR induced genes, however, where not similarly induced by TfoX production (e.g. *VCA0224)* or were also induced similarly in the qstR-minus strain in the previous data set (e.g. *VC1187,* 14.8-fold induction by TfoX, 14-fold in Δ*qstR* strain, Supplementary file 1). Since these genes do not appear to be QstR-dependent, they were not analyzed further.

**Figure 4.**
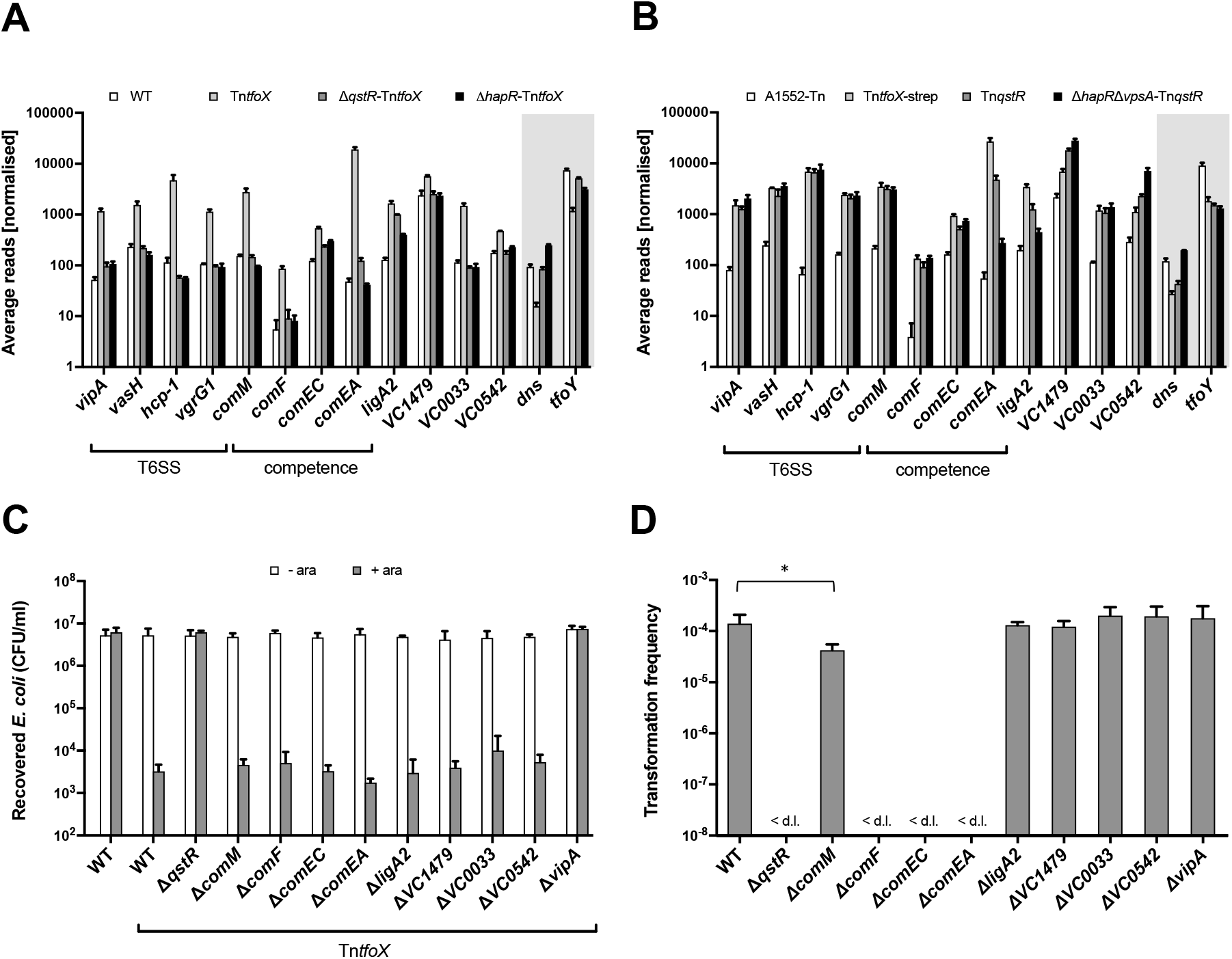
Contribution of QstR-regulated genes to transformation and interbacterial killing. **(A)** Average reads obtained from three RNA-seq experiments comparing expression of QstR-dependent genes in strain A1552 (WT), A1552-TntfoX, Δ*qstR-TntfoX* and Δ*hapR-TntfoX* (18,35). Shaded box indicates TfoX-repressed genes and the dependency of this repression on QstR and HapR. **(B)** Average reads obtained from three RNA-seq experiments comparing expression of QstR-dependent genes in strain A1552-Tn (carrying empty transposon), A1552-TntfoX-strep, A1552-TnqstR and Δ*hapR*Δ*vpsA-TnqstR* (Δ*hapR* strain that also carries a *vpsA* deletion to counteract enhanced biofilm production). Shaded box indicates the genes repressed in a QstR-dependent manner and the dependency of HapR in this repression. **(C)** Killing assay with tfoX-inducible strains and deletion of the indicated QstR-dependent genes. *E. coli* was used as prey and was co-cultured with each *V. cholerae* strain on LB agar plates without or with inducer (0.2% arabinose). The survival of the prey is represented as the number of colony forming units (CFU) per milliliter. **(D)** Transformation frequencies of strains with deletion of QstR-regulated genes assessed on chitin. For (C) and (D) the values shown are averages of three independent experiments with error bars representing the SD. Significant differences were determined by Student’s t-tests (* p<0.05, * * p<0.01, *** p<0.001; n.s., not significant).

A more detailed examination of the RNA-seq data revealed that QstR is sufficient to induce the expression of the majority of QstR-dependent genes (including all genes encoding proteins involved in T6SS) to a comparable extent as TfoX (Fig. 4B and S4A). This finding suggests that the role of TfoX in their regulation might be through TfoX’s ability to induce *qstR.* However, QstR was not sufficient to fully induce the expression of *comEA* since it was induced by QstR ˜ 87-fold compared to ˜490-fold by TfoX, in line with our previous report about *comEA* expression (34). The difference in *comEA* production between strains induced for TfoX or QstR individually was also observable at the protein level (compare lane 3 and 5 in Fig. S4B). However, co-production of both TfoX and QstR results in enhanced expression of *comEA* and elevated protein levels (Fig. S4A and S4B, lane 9) demonstrating that both regulators are required for full *comEA* induction. Furthermore, deletion of *hapR* had no effect on QstR induced expression of almost all the genes tested, except for *comEA* and *ligA2.* The transcript levels of these genes were markedly reduced in the Δ*hapR-TnqstR* strain compared to the TnqstR strain (Fig. 4B, Fig. S4A), suggesting an additional role of HapR (other than *qstR* activation) in the regulation of these genes. 8 genes were also repressed at least 2-fold when QstR was produced alone (Supplementary file 2), including *dns* (but not in the *hapR* mutant, see above) and *tfoY* (Fig. 4B).

Next, we attempted to evaluate the contribution of the genes induced by QstR to the known QstR-mediated phenotypes, i.e. natural transformation and interbacterial killing. As shown in Fig. 4C the killing of *E. coli* was unaffected in strains deleted for any of the QstR-dependent genes outside of the T6SS clusters (*vipA,* the first gene in the large T6SS cluster is shown as a representative example). As shown previously, deletion of *comEA, comEC* or *comF* all abolished transformation (15) while deletion of *comM* resulted in only a marginal (5-fold) reduction in transformation (Fig. 4D). Deletion of the other genes (i.e. *ligA2, VC1479, VC0033* and *VC0542*) did not have any apparent effects on transformability or interbacterial killing. The reason for their QstR-dependent regulation therefore remains unknown.

### QstR is sufficient to induce interbacterial killing but not competence

In line with the expression profiles of T6SS genes, a strain producing QstR from the transposon without any *tfoX* induction killed *E. coli* as efficiently as a strain induced for *tfoX* (Fig. 5A). In contrast, a strain producing the non-dimerizing QstR[L137A] variant was unable to kill *E. coli* (Fig. 5A). These results show that TfoX, once QstR expression is turned on, is no longer required and that QstR alone is sufficient for the induction of the T6SS, leading to its full functionality in interbacterial killing. Similarly, deletion of *hapR* in a QstR overproducing strain (i.e. Δ*hapR-TnqstR*) did not affect killing. However, this phenotype remained VasH-dependent (Fig. 5A). VasH, encoded within the major T6SS cluster, is a σ^54^ activator protein (19) that regulates transcription of the genes in the major and auxiliary T6SS clusters and is necessary for secretion (45,46). The requirement for VasH in the QstR-dependent T6SS activation is in line with a previous report showing that VasH is still necessary for TfoX-induced interbacterial killing (35).

**Figure 5.**
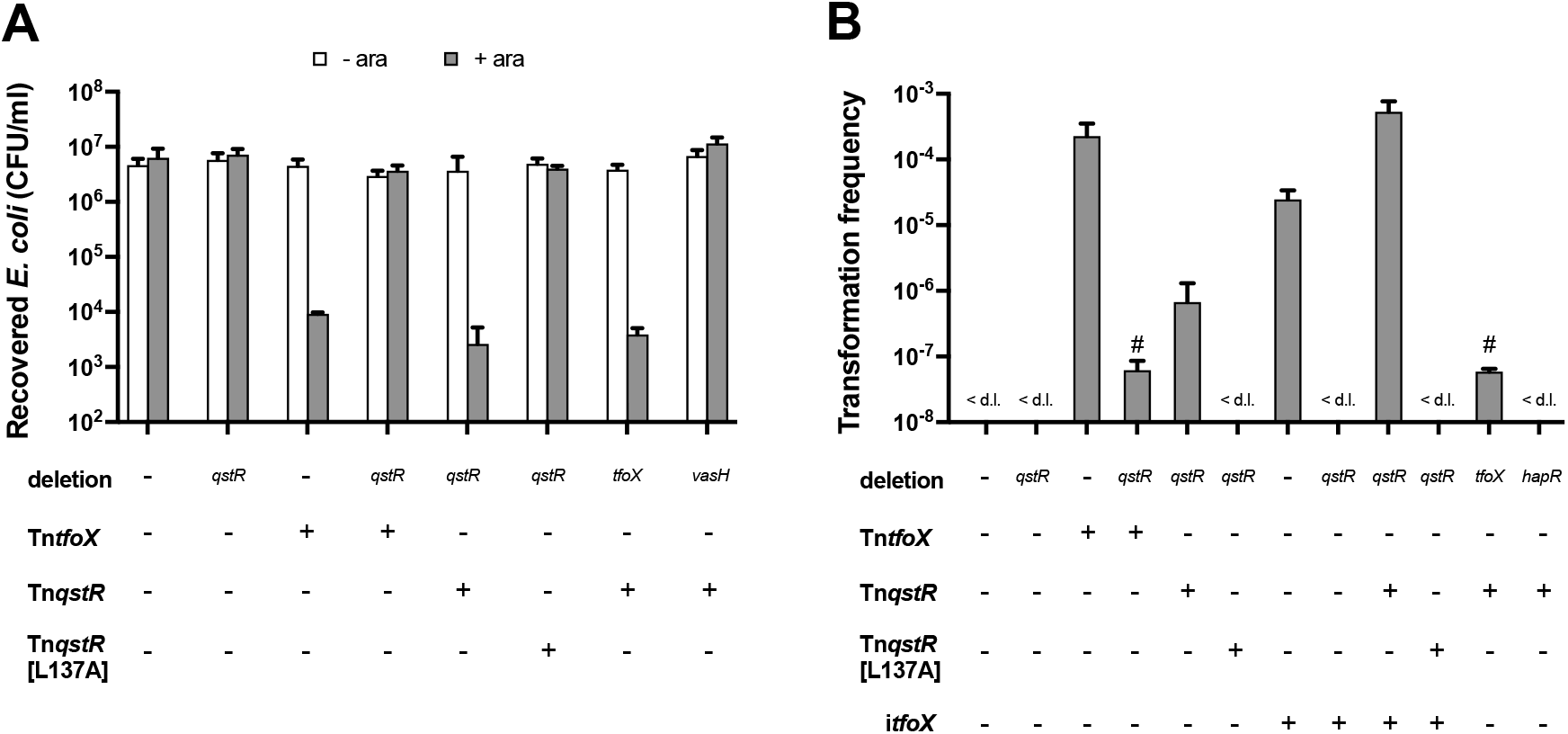
QstR production is sufficient to induce *E. coli* killing but not transformation. Killing assay **(A)** and chitin-independent transformation assay **(B)** of strains either without (−) or with (+) a transposon (Tn) with inducible copies of *tfoX, qstR* or the qstR[L137A] variant. Due to the insufficiency of *qstR* induction for transformation, a co-induction of *tfoX* was also tested (itfoX). The killing assay was performed as described in Fig. 4. Arabinose was used to induce the gene carried on the transposon and/or *itfoX.* In both panels the values are averages of three independent experiments with error bars representing the SD. <d.l., below detection limit; #, transformation frequency below the detection limit in two out of three experiments; the detection limit was therefore used for calculation of the average for this strains.

Contrary to T6SS activation, QstR alone was not sufficient for full competence induction. We did observe rare transformants upon TnqstR induction; however, the transformation frequency of a Δ*tfoX-TnqstR* strain was close to or below the detection limit (Fig. 5B). The inability of QstR to induce competence was not unexpected since the genes encoding components of the type IV pilus that forms the DNA-uptake machinery required for transformation (15) are activated by TfoX in a QS-independent and hence, QstR-independent manner (9,17). We therefore tested the effect of simultaneous induction of *tfoX* and *qstR* using a TnqstR-carrying strain that, in addition, carried an inducible copy of *tfoX* in the neutral *lacZ* locus (referred to as *itfoX*). This strain resulted in a significantly higher transformation frequency (Fig. 5B) and *comEA* expression (Fig. S4A and B) than the strains carrying either *itfoX* or TnqstR alone). These results demonstrate that both regulators are required for competence development.

### QstR directly regulates genes in its regulon by binding to DNA

To investigate whether QstR directly regulates target genes by binding to their promoters we first attempted electrophoretic mobility shift assay (EMSA) with purified QstR. We used probes encompassing the upstream regions of genes that are known to be affected by QstR production (Fig. 4A and B), namely *comEA, comM,* and *dns.* However, all attempts to observe specific QstR binding to these probes were unsuccessful. We speculated that the interaction with another protein or a cofactor might be required for QstR’s DNA binding ability and therefore repeated the experiment in the presence of *V. cholerae* cells lysates (e.g., of cells induced for competence and overproducing either QstR or its QstR[L137] variant). Still, no DNA binding of QstR occurred under those conditions. Hence, we decided to map the binding sites of QstR by chromatin immunoprecipitation coupled with deep sequencing (ChIP-seq). For this purpose we used cross-linked lysates of WT and Δ*qstR* cells grown under competence-inducing conditions. Sequencing of the DNA obtained by immunoprecipitation with a specific anti-QstR antibody revealed 52 QstR-dependent peaks. Importantly, we identified a peak upstream of most of the genes shown to be QstR-dependent, namely *comM, dns, comF, comEC, vipA* (the first gene of the large T6SS cluster), *ligA2, tfoY* and others (Fig.6A; ChIP DNA coverage Fig. S5, zoomed in peaks Fig. S6; full list of peaks in Table S2).

**Figure 6.**
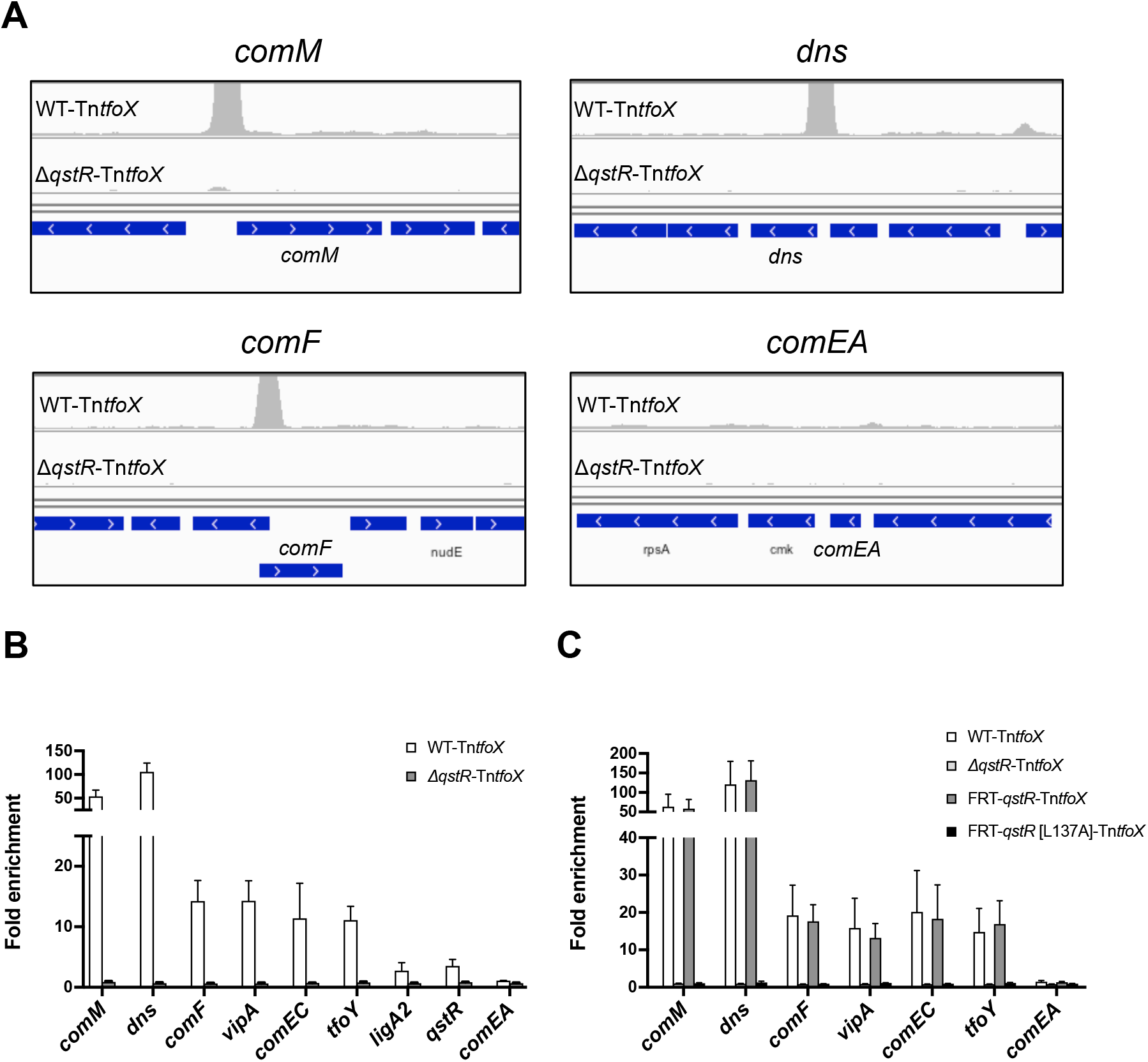
QstR binds to DNA upstream of regulated genes but not upstream of *comEA*. **(A)** Coverage of DNA reads for DNA obtained by chromatin immunoprecipitation with anti-QstR antibody coupled with sequencing (ChIP-seq) for the strains A1552-TntfoX (WT-TntfoX) and Δ*qstR-TntfoX.* The panels show DNA enrichment upstream of QstR-dependent genes *comM, dns* and *comF* but not *comEA.* The scale of the y-axis is set to the same level for all panels (0-50,000). The images were prepared using IGV software (63). **(B)** Enrichment of DNA fragments upstream of QstR-dependent genes determined by quantitative PCR (qPCR) using ChIP DNA. The fold enrichment was determined by dividing ‘Input %’ of the target by ‘Input %’ of the background (fragment upstream of *gyrA;* not enriched in the ChIP-seq experiment; see Fig. S5). **(C)** The non-dimerizing QstR mutant does not bind to DNA. The graph shows the fold enrichment of DNA fragments determined by ChIP-qPCR (as in panel B) for the strains *WT-TntfoX*, Δ*qstR-TntfoX, FRT-qstR-TntfoX* and FRT-*qstR[L137A]-TntfoX*. The inducer (arabinose) concentration was optimized to ensure comparable amount of QstR protein produced in each strain (0.02% ara for WT-TntfoX and Δ*qstR-TntfoX,* 0.01% ara for *FRT-qstR-TntfoX*, and 0.2% ara for FRT-*qstR[L137A]-TntfoX* (see Western blot Fig. S9). The values shown in (B) and (C) are averages from three independent experiments with error bars representing the SD.

Interestingly, we identified a QstR-dependent peak upstream of *VC0047* (Fig. S6, and Table S2), the first gene of a four-gene operon. The protein encoded by *VC0047* shares homology with TsaP of *Neisseria gonorrhoeae,* a peptidoglycan binding protein that forms a peripheral structure outside of the channel formed by the secretin PilQ and is important for type IV pilus biogenesis (47). Deletion of *VC0047* in *V. cholerae* reduced the frequency of transformation ˜10-fold (transformation frequency on chitin 1.4 × 10^−5^ [± 8.02 × 10^−6^] compared to the WT parent 1.4 × 10^−4^ [± 6.93 × 10^−5^]). The second gene in the operon encodes DprA, a homolog of a single stranded DNA binding protein that is required for transformation in *V. cholerae* (9) and other bacteria (48–50). Both genes are induced by TfoX but their induction appears to be QS-independent ((11); Supplementary file 1). Nevertheless, production of QstR from TnqstR induced the expression of *VC0047* ˜2-fold as determined by RNA-seq (Supplementary file 2) and 5-fold in the qRT-PCR experiment (Fig. S4A). Additionally, co-induction from *itfoX* and TnqstR resulted in considerably higher *VC0047* mRNA levels than when *tfoX* alone was induced (Fig. S4A). Therefore, it appears that QstR may play an auxiliary role in the regulation of this promoter.

Another gene that is induced during competence in other bacteria is *radC* (51–54). *radC* encodes a DNA repair protein that appears to be dispensable for transformation in *S. pneumoniae* and *H. influenzae* (55,56). RadC also appears dispensable for transformation in *V. cholerae* (transformation frequency on chitin is 2.49 × 10^−4^ [± 5.22 × 10^−5^] compared to the one of the WT, which is 1.4 × 10^−4^ [± 6.93 × 10^−5^]). We identified two QstR-dependent peaks in the proximity of this gene (Fig. S6; Table S2). One peak was located in the beginning of the preceding gene (*VC0216;* 1783 bp upstream of *radC*) and the second one was 118 bp upstream of *radC.* However, the expression of *radC* was not affected at significant levels upon TfoX or QstR induction (Supplementary file 1 and 2). Surprisingly, we also found a peak upstream of *pilA,* which encodes the major subunit of the DNA-uptake pilus (Fig. S6).

We previously showed that *pilA* expression is highly induced by TfoX in a QS-independent manner (11) and confirmed this result here, by qRT-PCR (Fig. S4A). Therefore it is unclear if and how QstR binding to these loci might affect the expression of these genes.

Furthermore, we identified a peak upstream of *qstR* itself (Fig. S6) suggesting that *qstR* may be subject to auto-regulation. Indeed, using discriminatory primers that either anneal to the native *qstR* locus or to *qstR* inside the TnqstR construct, we could demonstrate that QstR induces its own expression, though to a lesser extent in the absence of TfoX (Fig. S7). Moreover, as shown in Fig. S7, auto-regulation of QstR was fully dependent on the presence of HapR, in agreement with recent observations using a plasmid-borne system based on a luciferase reporter in strain *V. cholerae* C6706 (57).

To validate the ChIP-seq data we performed ChIP-qPCR experiments using primers amplifying the regions around the identified peaks. These experiments confirmed the enrichment of the top 15 peaks identified by ChIP-seq, as well as those for *pilA, qstR,* and *VC0047* (Fig. 6B and S8). Additionally, using ChIP-qPCR we showed that the non-dimerizing QstR[L137A] mutant does not bind to DNA, suggesting that dimerization is required for DNA binding (Fig. 6C and S9).

### QstR does not bind upstream of *comEA*

To our surprise we did not identify a QstR-dependent peak upstream of *comEA* in the ChIP-seq experiment and likewise, when validated by ChIP-qPCR, we did not observe any enrichment of DNA fragments in this region (Fig. 6A and B). This was unexpected since *comEA* is one of the most upregulated genes during competence induction and this upregulation is strictly QstR-dependent (17,18) as also demonstrated in this study (˜ 400-fold induction in *TntfoX* strain vs. ˜ 2-fold in Δ*qstR-TntfoX* strain; Supplementary file 1 and Fig. S4A). We therefore hypothesized that *comEA* expression might be controlled by another regulator that is produced in a QstR-dependent manner. In an attempt to identify such a regulator we first deleted the non-essential genes in proximity of the ChIP-seq peaks that could potentially be regulated by QstR and tested *comEA* expression using a chromosomal *comEA-lacZ* transcriptional reporter fusion. However, none of these genes altered *comEA-lacZ* expression significantly, either in the presence or absence of TfoX (Fig. S10). Additionally we tested these deletion strains for their natural transformability and *E. coli* killing and discovered that none of these genes was required for transformation (Fig. S11A) or T6SS-mediated interbacterial killing (Fig. S11B). Deletion of *VC2320* abolished the killing phenotype; however, the strain had a significant growth defect. We also observed impaired killing by the Δ*dksA* strain. *dksA* encodes a global regulator involved in the stringent response and its deletion was previously shown to affect processes such as motility and virulence in *V. cholerae* (58). Therefore, none of these genes appears to encode proteins specifically involved in transformation or killing.

Next, we attempted to identify the putative intermediate regulator using a transposon mutagenesis screen. Using a strain carrying the *comEA-lacZ* transcriptional reporter fusion and TntfoX, we screened ˜ 80,000 colonies (in two separate screens) for loss of *lacZ* expression. As expected, we identified multiple transposon insertions in the genes encoding known *comEA* regulators such as *qstR, hapR, cyaA* and *cytR,* as well as in several other genes (Table S3; details are provided in the supplementary material). However, none of these newly identified genes encoded a specific QstR-dependent regulator of *comEA.*

As a final approach to identify the potential regulatory protein, we directly determined the proteins that bind to the *comEA* promoter region. To do so, we performed a DNA affinity pull-down assay (41,59) using a biotinylated probe encompassing the upstream region of *comEA.* This probe was incubated with the lysates of competence-induced WT or qstR-minus strains. As a negative control, we used the upstream region of *gyrA.* To validate the assay, we used a probe containing the upstream region of *qstR* incubated with the same lysates, and confirmed HapR’s binding to this region (Fig. S12A). Mass spectrometry analysis of the eluted fractions revealed several DNA binding proteins that were common to all the probes. Additionally, a few proteins were specifically eluted from the *comEA* probe. Most notably VCA0199 (uncharacterized protein) was highly enriched on the *comEA* probe, as well as two other transcriptional regulators (Table S4). However, strains carrying deletions of these genes were fully transformable and *comEA-lacZ* expression appeared unaffected (Fig. S12 B and C). Importantly, we did not observe any significant differences in the elution profiles from the *comEA* probe incubated with the lysates of either the wild-type or Δ*qstR* strains. In conclusion, despite several attempts, we were so far unable to identify the putative intermediate regulator of *comEA.*

## Discussion

QstR has been shown to be an important regulator involved in natural competence and TfoX-dependent activation of the T6SS genes, though the mode of action and the full set of regulated genes was unknown (17,18). Here, we show that QstR undergoes dimerization and that dimerization is required for binding to DNA upstream of QstR-regulated genes. Dimerization and the activity of LuxR-type regulators are often controlled by phosphorylation (42). However, similar to the biofilm regulator VpsT, the putative phosphorylation site of QstR appears dispensable for protein function. Interestingly, in LuxR-type regulators that act independently of phosphorylation, a co-factor is often required to mediate dimerization and hence DNA-binding (23–25). In the case of VpsT this cofactor is the nucleotide second messenger c-di-GMP (26). We therefore previously hypothesized that QstR activation might require the binding of a cofactor, perhaps competence specific (17). However, for the reasons that follow, we no longer favor the idea that such a cofactor would act in a similar manner as c-di-GMP in VpsT, which primarily fosters oligomerization (26).

First, QstR is able to self-interact in an *E. coli* bacterial two-hybrid assay. Second, QstR purified from *E. coli* behaves as a dimer in solution, as evidenced by analytical SEC. Thus, these results demonstrate that QstR can dimerize in a heterologous system without any *Vibrio-specific* signals. Third, in contrast to VpsT, QstR contains a proline substitution in the third position of the conserved c-di-GMP binding motif (i.e. WLPR), suggesting that QstR is unlikely to bind c-di-GMP. Indeed, our results demonstrate that QstR is highly functional under low c-di-GMP conditions but that a variant with the restored canonical binding motif (i.e. WLTR) is rendered c-di-GMP dependent. Moreover, substitution of the conserved arginine within the motif, required for c-di-GMP binding in VpsT, does not abolish QstR function. These findings are in agreement with a recent report on the VpsT homolog of *Vibrio vulnificus,* BrpT, which bears the same WLPR variant motif and also acts in a c-di-GMP independent manner (60). Although we cannot exclude the possibility that another cofactor might be required for QstR dimerization and full functionality, it appears that it would not be competence-or even species-specific. Finally, multiple lines of evidence demonstrate that the L137A substitution abolishes QstR function by rendering it unable to dimerize. Thus, in contrast to VpsT, which contains a second nucleotide-independent dimerization domain (26), QstR likely contains a single dimerization interface.

The regulation of *comEA, comEC, dns,* and the T6SS genes was previously shown to be QstR-dependent (17,18). Using RNA-seq we defined the QstR regulon and identified other genes that are regulated in a QstR-dependent manner (Fig. 7). None of the QstR induced genes, besides the known clusters, affected T6SS-mediated interbacterial killing. However, two genes involved in transformation were identified as being QstR-dependent, namely *comF,* which was previously shown to be essential for transformation due to its role in translocation of DNA across the inner membrane (15) and *comM.* In a recent study, *comM* was shown to encode a helicase that promotes integration of the incoming heterologous DNA into the genome (61), similar to RadA’s role in natural transformation of *S. pneumoniae* (62). Moreover, deletion of *comM* reduced transformation frequencies ˜ 100-fold (61). Here, however, the defect of a comM-minus strain was only modest with a ˜ 5-fold reduced transformation frequency. The reason for this discrepancy is most likely based on the different transforming DNA used in both assays. We used genomic DNA with an integrated antibiotic resistance cassette, a DNA substrate that likely resembles DNA released during the co-regulated interbacterial killing events that occur on chitinous surfaces (18), while Nero *et al.* used a PCR product with shorter homologous regions. Additionally, QstR might also play an auxiliary role in regulation of other genes encoding proteins involved in competence such as *VC0047* and *dprA* (*VC0048*). The remaining newly identified QstR-induced genes did not appear to be required for either of the two known QstR-dependent phenotypes and thus, the significance of their regulation by QstR remains unknown.

**Figure 7.**
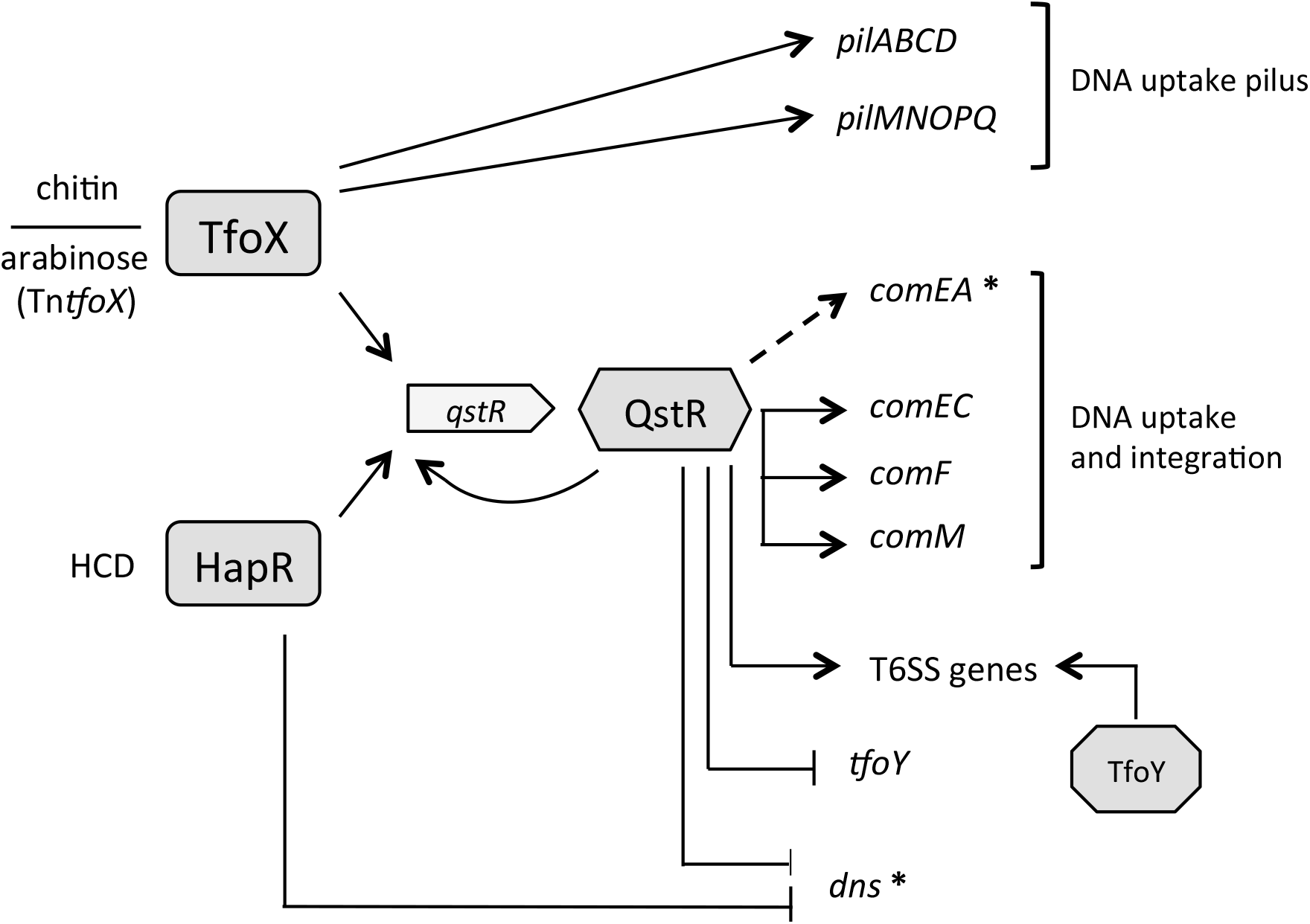
Model of the QstR-dependent regulation of competence and type VI secretion. Chitin degradation compounds induce production of the master regulator TfoX, which then activates expression of the genes encoding the components of the DNA uptake pilus. In the chitin-independent system expression of *tfoX* from the transposon (*TntfoX*) is induced by arabinose. Both TfoX and the quorum sensing regulator HapR, that accumulates at high cell density (HCD), are required for production of QstR. QstR, which induces its own transcription, activates expression of the genes involved in DNA uptake and integration, and also type VI secretion (T6SS major cluster) by binding to their promoter regions. Additionally, QstR directly represses transcription of *dns* (encoding nuclease) and *tfoY* (encoding another regulator of T6SS).*, regulation of these genes by QstR also requires HapR. Dashed line: regulation of *comEA* by QstR is likely achieved via an unknown intermediate regulator and also requires TfoX.

Notably, when we mapped QstR binding sites by chromatin immunoprecipitation coupled with deep sequencing (ChIP-seq), we found that QstR binds upstream of the majority of the QstR regulated genes, including its own promoter region, suggesting that transcriptional regulation by QstR is direct. Interestingly, since the expression of *dns* (nuclease) and *tfoY* (T6SS regulator) are both reduced in a QstR-dependent manner, yet we observed QstR-dependent enrichment of DNA fragments upstream of these genes, we conclude that QstR is a dual regulator that can act as a transcriptional activator and repressor. However, analysis of the DNA sequences upstream of the genes identified by ChIP-seq did not reveal an obvious common binding motif. Further investigation into the nature and positioning of the QstR binding sites will therefore be required to clarify how the binding site determines whether the gene will be up-or down-regulated.

Our results also demonstrate that QstR is itself sufficient for the expression of the majority of QstR-dependent genes, including all genes encoding the T6SS components. Accordingly, TfoX-independent induction of QstR production is sufficient for activation of T6SS mediated interbacterial killing, corroborating a previous report in a different *V. cholerae* strain (57). Thus, the role of TfoX in chitin-dependent T6SS activation appears to be through its primary role in *qstR* induction.

In contrast, TfoX is required for the regulation of the majority of the genes involved in transformation, in particular ones that encode the components of DNA-uptake machinery, including the QstR-dependent *comEA* gene.

Unexpectedly, we did not find evidence that QstR binds directly to the *comEA* promoter despite it being the most highly induced gene during competence development ((3,11,17,18) and this work). One possibility is that this reflects a technical artifact of the ChIP or ChIP-qPCR methods. However, the fact that it was possible to identify the other genes that are induced or repressed at varying levels in a QstR-dependent manner, argues against this idea. Our results therefore strongly suggest that QstR does not regulate *comEA* directly, but rather requires the involvement of an intermediate regulator. In support of this hypothesis is the observation that in contrast to majority of the QstR-dependent genes, HapR is required for the QstR mediated activation of *comEA* in the absence of *tfoX* induction. Yet, HapR did not bind to the *comEA* promoter region, either in previous EMSA experiments (17) or in the DNA affinity pull-down experiment presented here (Fig. S12A), whereas it bound to the *qstR* probe in both assays. Thus, it seems plausible that this putative regulator could be dependent on both QstR and HapR. Nevertheless, despite using various complementary approaches we were unable to identify such a regulator. Future work should therefore be directed towards finding the missing regulator, which might provide new insights into the complex regulatory pathways that lead to competence development in *V. cholerae*.

## Data availability

The RNA-seq data are accessible through the GEO Series accession number GSE114592.

## Acknowledgments

We thank Stefanie Boy-Röttger for her assistance with the size exclusion chromatography and Romain Hamelin and the whole team of the EPFL Proteomics Core Facility for the mass spectrometry analysis of protein samples. We also thank Fitnat Yildiz for plasmid pSC189, Marie de Barsy for the ChIP protocol, Julia van Kessel for kindly providing the DNA affinity pull-down protocol and David Adams for comments on the manuscript.

